# Development of a novel HPTLC-based method for the simultaneous quantification of clinically relevant lipids from cells and tissue extracts

**DOI:** 10.1101/779603

**Authors:** Michelle Pinault, Cyrille Guimaraes, Céline Ben Hassen, Jorge L. Gutierrez-Pajares, Stéphan Chevalier, Caroline Goupille, Pierre Bernard-Savary, Philippe G. Frank

**Affiliations:** Université de Tours, INSERM, UMR1069, Nutrition, Croissance et Cancer, Tours, France; CHRU Hôpital Bretonneau, Tours; Chromacim, 170 Rue de Chatagnon, 38430 Moirans

**Keywords:** Cholesterol, Triglycerides, Fatty Acids, Cancer

## Abstract

Lipids such as cholesterol, triglycerides, and fatty acids play important roles in the regulation of cellular metabolism and cellular signaling pathways and, as a consequence, in the development of various diseases. It is therefore important to understand how their metabolism is regulated to better define the components involved in the development of various human diseases. In the present work, we described the development and validation of an HPTLC method allowing the separation and quantification of free cholesterol, cholesteryl esters, non-esterified fatty acids, and triglycerides. This method will be of interest as the quantification of these lipids in one single assay is difficult to perform.

## Introduction

Abnormality in the metabolism of lipids such as cholesterol (free or esterified), non-esterified fatty acids, and triglycerides have been shown to control the development of various diseases including atherosclerosis, obesity, cancer, nonalcoholic fatty liver disease, as well as other metabolic diseases. [1–4]. In cancer, they have been demonstrated to play an important role in the regulation of cancer progression. Their functions include the regulation of cellular signaling pathways involved in cell cycle regulation as well as other pathways regulating cancer cell migration and survival [5–7]. In several types of cancer, plasma cholesterol levels have been shown to regulate tumor formation and cancer aggressiveness [8, 9]. In vascular disease, studies have now demonstrated the role of cholesterol and triglycerides in the development of atherosclerosis [10]. After accumulation in the form of LDL-derived cholesterol in the arterial wall of blood vessels, an inflammatory reaction leads to the attraction of monocyte/macrophages in the intima and eventually atheroma development [11]. Finally, obesity, which is characterized by increased adipose tissue and increased triglyceride accumulation, is associated with defect in the metabolism of many lipids such as cholesterol and fatty acids, and its appearance is associated with the development of other diseases such as cardiovascular disease, type 2 diabetes, and cancer [12].

Taken together, these data suggest that a method that could facilitate the simultaneous cellular or tissue quantification of these lipids and allow valid comparisons between samples would be invaluable to study the regulation and the role of these lipids in human diseases. In the present study, we described the development of a method for the reproducible quantification of these lipids (**Fig. 1**) in cells and tissue extracts.

**Fig. 1:**
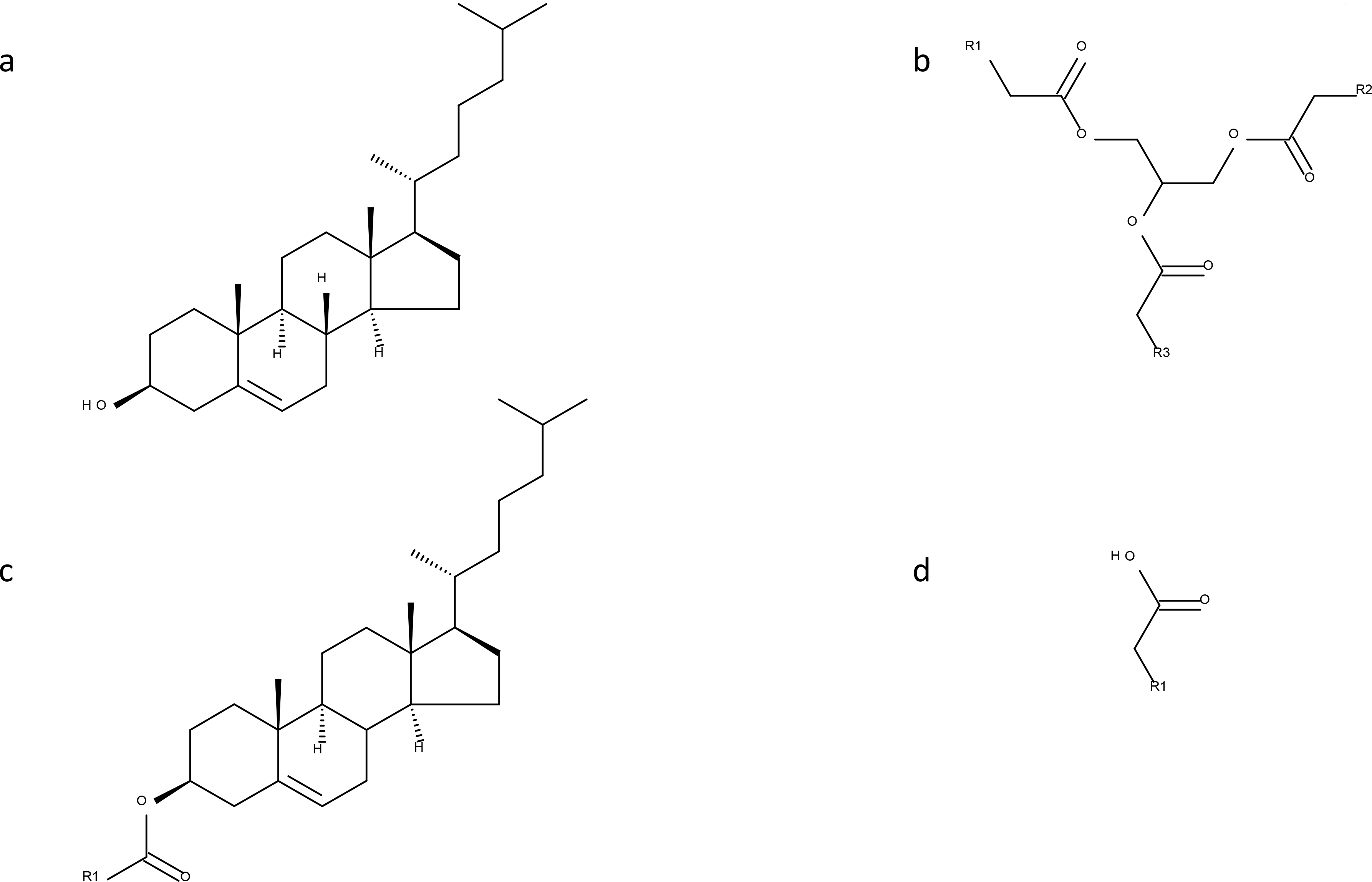
Lipid structure representations. a. Free cholesterol. b. Triglyceride. c. Cholesteryl ester. d. Non-esterified fatty acid. R1, R2, and R3 represent alkyl groups.

## Materials and methods

### Reagents

Free cholesterol (FC), cholesteryl oleate (CE), oleic acid (NEFA) and triolein (TG) were obtained from Sigma-Aldrich (Saint-Quentin Fallavier, France). All other reagents were analytical grade.

Phospho-vanillin was prepared as followed: 0.99 g of Vanillin (Sigma-Aldrich) was dissolved in 10 mL ethanol 100% before dilution into 153mL with distilled water. This solution was mixed with 337 mL of 85% H_3_PO_4_ (w/v) with constant stirring. The resulting solution was prepared before each use and kept in a dark bottle at room temperature. Perchloric acid (65%) was obtained from Sigma-Aldrich.

### HPTLC instruments and chromatographic conditions

Standard calibration curves were obtained by spotting standards of known lipid composition. For sample analyses, after lipid extraction with the method from Bligh and Dyer [13], lipid extracts were also spotted on HPTLC precoated silica gel glass plates 60F_254_ (20×10 cm; 1.05642.0001; E. Merck, Darmstadt, Germany) using a Camag autosampler ATS4 sample applicator (Camag, Muttenz Switzerland) equipped with a 25-μL dosing syringe. For each plate, 14 tracks were used, five were used for calibration standards and nine for sample application. In successive plates, calibration standards were spotted on different tracks to avoid side effects.

Before each experiment, plates were pre-developed once in chloroform-methanol 1:1 (v/v), air-dried, and activated at 110°C in an oven for 20 min. Samples were applied in the form of 10-mm band shapes, 1.5cm from the bottom of the Silica plate at a constant flow rate of 150 nL/s. under nitrogen flow. This spray-on technique and the flow rate adjusted to the type of sample solvent to be applied, dries out the solvent during the application. It avoids diffusion and permits an optimal sample band focus that allows a good separation and a homogenous quantification. For development, the mobile phase used was a mixture of hexane:ether:acetic acid (70:30:1; v/v/v). Linear ascending development was carried out in a 20×10cm twin trough glass chamber (Camag, Muttenz, Switzerland) pre-equilibrated with the mobile phase. Optimal chamber saturation time for the mobile phase was 15 min at room temperature without saturation paper. The length of development was 10 cm. We used 20 mL of mobile phase for saturation and migration (10 mL in the tank facing the plate and 10mL in the tank where the plate is developed). Subsequent to the development, HPTLC plates were air-dried under a ventilated hood for 90 min after migration. Dried plates were treated by dipping for 1 min in an immersion tank containing a 50% perchloric acid solution. After a 2-hour drying period at room temperature, plates were placed on a plate heater (TLC Plate Heater 3, Camag Muttenz Switzerland) for 16 min at 160°C to allow carbonization for the staining reaction. Densitometric analysis was performed using a TLC/HPTLC video-densitometer (TLC Visualizer 2 Camag, with Videoscan software, Camag, Muttenz, Switzerland).

### Standard solution preparation

To establish the assay range, the limit of detection was determined for all applications. We tested concentrations ranging from 0.04 mg/ml to 0.8 mg/ml. A standard solution at 1mg/mL was prepared and used for the calibration curve. For each authentic standard of lipid, we performed a calibration curve with 5 points at increasing levels to quantify lipids on the plates. Stock solutions and standards were aliquoted and kept in glass tubes (with a screw cap and silicone/PTFE seal) at −20 °C. Under these conditions, products were stable for one year.

### Method validation

The method was validated as described by the ICH harmonization guidelines [14] in terms of linearity, accuracy, specificity, repeatability, system repeatability, and intermediate precision of measurement for peak area.

The percent of recovery was calculated by using the formula, % recovery = [(T-A)/S]×100, where: T is the percentage added to the maximum value in range (i.e. 50, 100, 150, and 200%), S is the maximum value in range, A is the difference between the area at the considered percentage and the area of S. Recovery studies were carried out by spiking the stock solutions (1 mg/mL) with 50%, 100%, 150%, and 200% of the standard lipid solution. After dilutions were made, recovery studies were performed. Precision of the measurements was verified by repeatedly testing (n=10) each mixed standard solutions (five different concentrations) containing all the lipids. Experiments were repeated the same day using several HPTLC plates.

LOD and LOQ were calculated from the following equations: LOD=3×(SD/S) and LOQ=10×(SD/S), with S, slope of the calibration curve and SD, standard deviation of the blank signal (n=10 measurements on different plates). For this determination, mean blank was obtained for each lipid class using a blank lane by spotting the solvent (chloroform/methanol, 2:1, v/v) and reading the signal at the same retention factor (Rf) as that of the corresponding lipids.

The limit of quantification (LOQ) was also estimated using the Eurachem approach [15]. Five concentration close to the quantification point were used, and the relative standard deviation was plotted against the measured concentration. The corresponding limit of quantification was obtained from this graph for a relative standard deviation of 10%.

Statistical analyses were performed with StatPlus 6.03.

### Spectrophotometric quantification of lipids using phospho-vanillin

The sulfo-phospho-vanillin assay [16] was performed to determine the accuracy of our method. Briefly, two mL of concentrated H_2_SO_4_ (98% w/v solution) were added to standards at different concentrations after solvent evaporation. Tubes were heated for 20 min in a boiling water bath and cooled down for 4 min. To each tube, 4 mL of the phospho-vanillin reagent were added for color development. After a 15-minute cooling period at 25 °C, absorbance was read on a Secoman-S1000 spectrophotometer.

### Cell culture

The MCF-10A cell line was obtained from the American Tissue Culture Collection (ATCC) (Molsheim, France). The MDA-MB-231 and MCF-7 cell lines were obtained from Cell biolabs, Inc. (San Diego, CA) and GenTarget (San Diego, CA), respectively. MCF-10A cells were grown DMEM /F12 Ham’s Mixture supplemented with 5% horse serum (Sigma-Aldrich), EGF (20 ng/ml, Sigma-Aldrich), insulin (10μg/ml, Sigma- Aldrich), hydrocortisone (0.5 mg/ml, Sigma- Aldrich), cholera toxin (100 ng/ml, Sigma-Aldrich), 100 units/ml penicillin and 100 μg/ml streptomycin. The other cell lines were grown in Dulbecco’s modified Eagle’s media (DMEM) containing 10% Fetal Bovine Serum (FBS) and 100 units/ml penicillin and 100 μg/ml streptomycin. Cells were placed in a humidified incubator kept at 37°C with 5% CO_2_. For lipid extraction, cells were grown in T175 flash to 90-95% confluence and were washed with PBS. They were harvested using a cell scraper. Lipids from the cell pellets were extracted using the method Bligh and Dyer [13].

For fluorescent microscopy, cells were seeded on coverslips and incubated in complete media. After 1 day in culture, cells were fixed in 2% paraformaldehyde, incubated with 1% BSA in PBS, and incubated with 300 nM Nile red (in PBS containing calcium and magnesium (CM)). After several wash in PBS-CM cells were mounted using SlowFade® diamond antifade reagent containing the nuclear stain DAPI (Fischer Scientific). Cells were observed on a Nikon TI-S microscope and analyzed using both NIS-BR software (Nikon, France).

## Results and Discussion

The method presented in this study was directly adapted from a TLC method previously used in our laboratory to separate fatty acids from triglycerides [17]. We tested it to determine whether we could simultaneously separate, detect, and quantify free cholesterol, non-esterified fatty acids, triglycerides, and cholesteryl esters (**Fig. 1**).

### Calibration curves – Linearity and range

A mixture of chloroform and methanol (2:1, v/v) was used to prepare solutions for all lipids. Using an automated sample applicator, spots were made for each range of mixtures containing free cholesterol, non-esterified fatty acids, triglycerides, and cholesteryl esters at increasing concentrations. A range of concentrations (each one repeated 10 times) was loaded onto each plate. Plates after run and the resulting chromatograms (example presented **Fig. 2**) allowed us to obtain the calibration curves presented in **Fig. 3**. All lipids were well separated and formed identifiable bands.

**Fig. 2:**
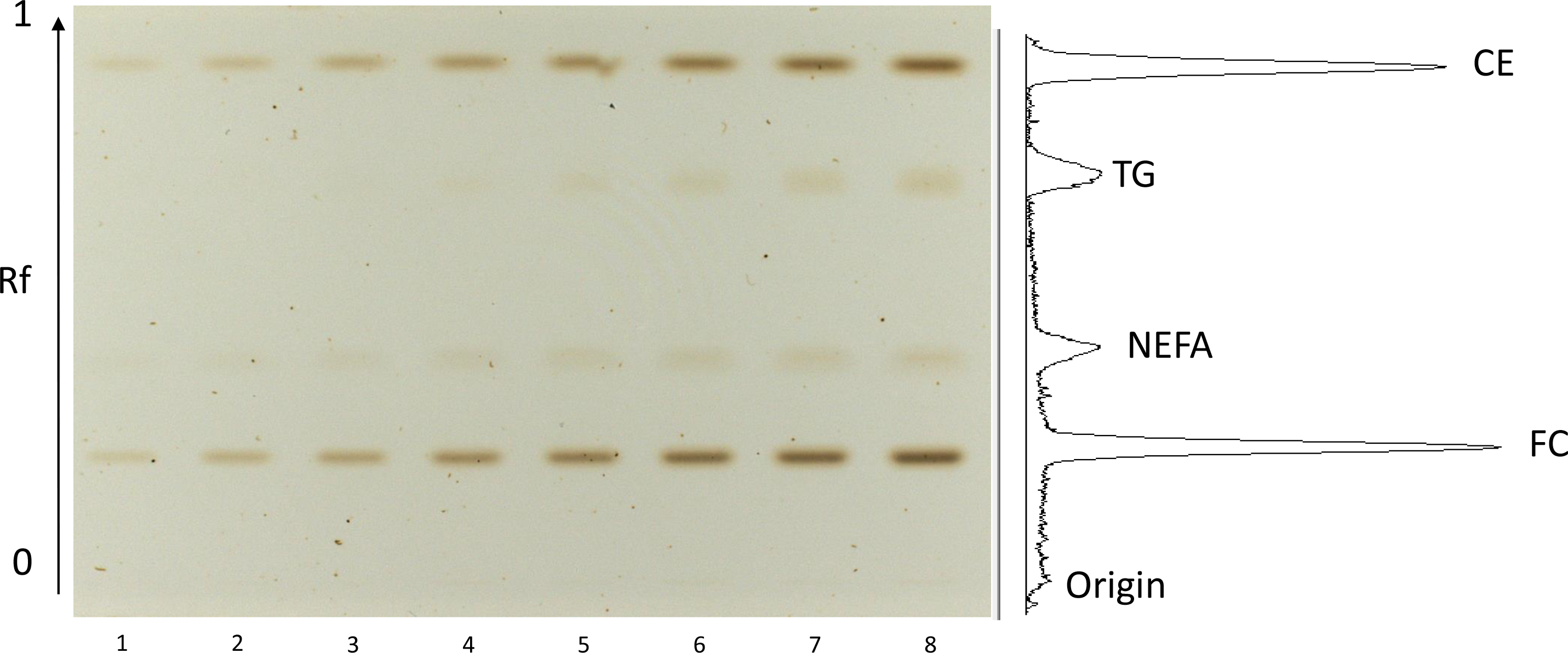
Profile of a standard separation realized by HPTLC. Samples, in the form of 7-mm bands, were spotted at a constant application rate of 150 nL/s under nitrogen (6 bars). For development, the mobile phase used was a mixture of hexane:ether:acetic acid (70:30:1; v/v/v). Linear ascending development was carried out in a 20×10 cm twin trough glass chamber (Camag, Muttenz, Switzerland) pre-equilibrated with the mobile phase. Optimal chamber saturation time for the mobile phase was 15 min at room temperature. The length of development was 7 cm. Subsequent to the development, HPTLC plates were air-dried and then treated by immersion in a 50% perchloric acid solution for 2 min. After a 2-hour drying period, plates were placed on a plate heater to allow carbonization. Lane 1 to 9: standards curves of each class of lipids. Amounts of standard loaded (lane number: μg): 1: 1; 2: 2; 3: 3; 4: 4; 5: 5; 6: 6; 7: 7; 8: 8. Images of standards were transferred to the Videoscan software, and densitometric scan were obtained. The densitometric scan presented in this figure was obtained for the 8μg standards. FC: free cholesterol, NEFA: non-esterified fatty acids, TG: triglycerides, CE: cholesteryl esters.

**Fig. 3:**
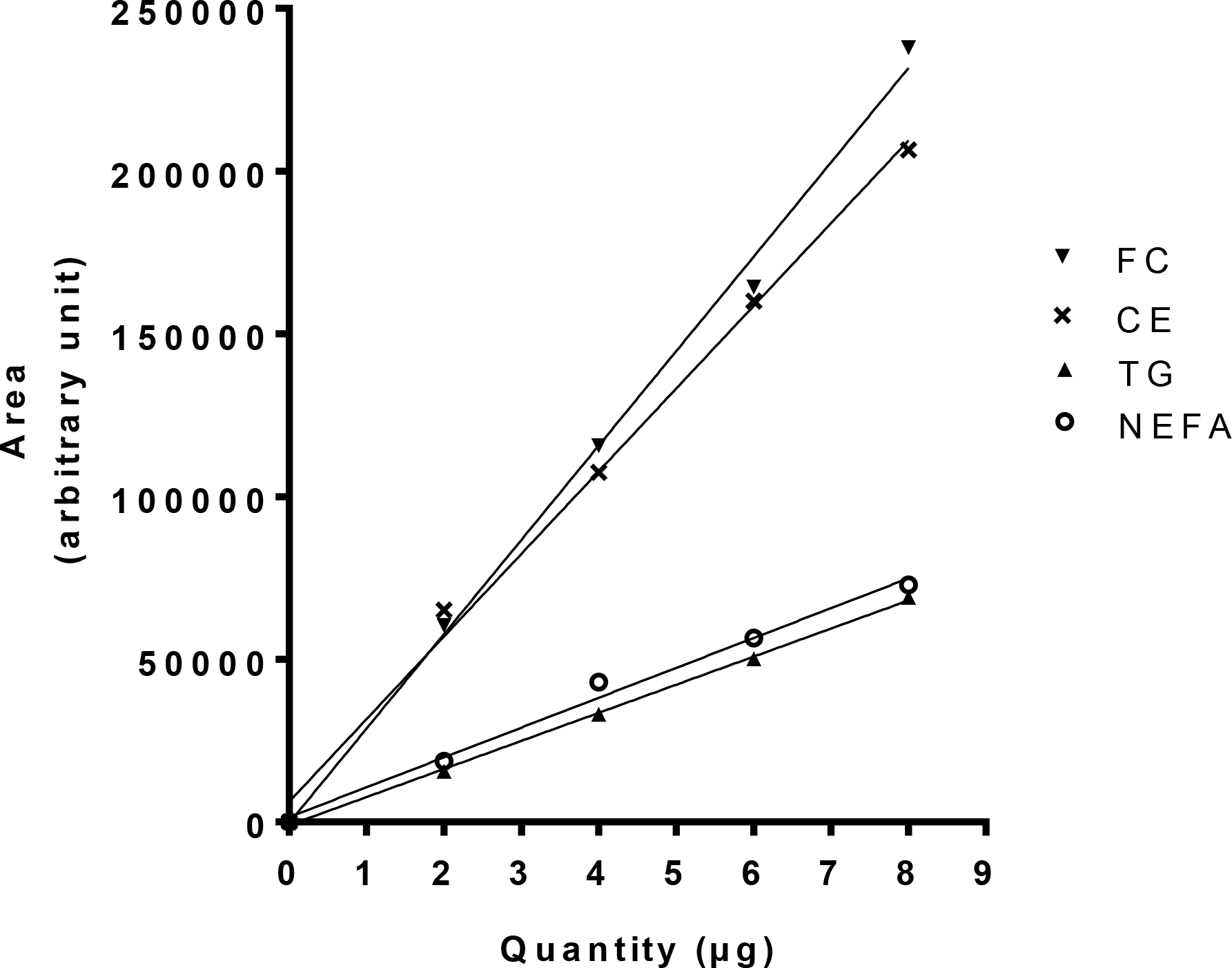
Calibration curves of the different lipids separated by HPTLC. Using an automated sample applicator, deposits were made for each range of mixtures containing free cholesterol, non-esterified fatty acids, triglycerides, and cholesteryl esters at increasing concentrations. Images of standards and samples were transferred to the Videoscan software (version 1.2), and calibration curves were obtained by plotting the peak area under the curve against the concentration of applied standards. Standard curves were obtained for all the analyzed lipids. FC: free cholesterol, NEFA: non-esterified fatty acids, TG: triglycerides, CE: cholesteryl esters.

Images of plates with standards were transferred to the Videoscan software. After densitometric analysis, calibration curves were obtained by plotting the peak area under the curve against the concentration of standards. Standard curves were obtained for each of the analyzed lipids. **Table 1** shows Rf values obtained for free cholesterol, non-esterified fatty acids, triglycerides and cholesteryl esters. In all cases, linearity was observed at levels ranging from:

- 1000 to 8000 ng/spot for free cholesterol (FC).
- 2000 to 8000 ng/spot for non-esterified fatty acids (NEFA).
- 2000 to 8000 ng/spot for triglycerides (TG).
- 1000 to 8000 ng/spot for cholesteryl esters (CE).

**Table 1:** HPTLC profile of a standard lipid solution.

| Lipid | Peak Beginning |  | Peak Maximum |  |  | Peak End |  | Peak Area |  |
| --- | --- | --- | --- | --- | --- | --- | --- | --- | --- |
|  | Rf | H | Rf | H | [%] | Rf | H | A | [%] |
| CE | 0.828 | 0.0 | 0.881 | 2121.8 | 30.05 | 0.897 | 0.0 | 24292.9 | 28.81 |
| TG | 0.769 | 0.0 | 0.800 | 646.1 | 9.15 | 0.825 | 9.8 | 7998.5 | 9.49 |
| NEFA | 0.359 | 26.1 | 0.413 | 506.8 | 7.18 | 0.473 | 19.1 | 9861.1 | 11.69 |
| FC | 0.179 | 210.8 | 0.203 | 3786.8 | 53.63 | 0.228 | 293.8 | 42168.7 | 50.01 |

Rf: retention factor, ratio distance migrated / total distance covered by the solvent.
H: peak height.
These values were obtained with 8 µg/spot for each lipid.
FC: free cholesterol, NEFA: non-esterified fatty acids, TG: triglycerides, CE: cholesteryl esters.

Correlation coefficients (*r*^2^) are presented in **Table 2** for the calibration curves shown in **Fig.3**. Linearity was confirmed using 10 measurements per concentration. Fives different plates were used.

**Table 2:** Linear regression data obtained for the different calibration curves.

| **Lipid** | **Slope (± *RSD*)** | **Correlation coefficient** | **Equation of the regression line**<br>**(peak surface =slope x amount deposited)** |
| --- | --- | --- | --- |
| FC | 28590 (± 3622) | 0.9959 | $y = 28590x + 1605$ |
| NEFA | 9182 (± 1284) | 0.9952 | $y = 9182x + 49$ |
| TG | 7969 (± 1371) | 0.9933 | $y = 7969x - 1300$ |
| CE | 26657 (± 4621) | 0.9940 | $y = 26657x + 510$ |

FC: free cholesterol, NEFA: non-esterified fatty acids, TG: triglycerides, CE: cholesteryl esters.

### Accuracy of the Method

We compared our technique with the method of Knight *et al*, [16] originally used to quantify plasma lipids levels. This method was first adapted for oleic acid and compared to the original work of Knight *et al.* (**Fig. 4**). A small relative deviation of 1.6%, was observed. In a separate experiment, concentration ranges of triglycerides, non-esterified fatty acids, free cholesterol and cholesteryl esters were also analyzed with this spectrophotometric method (**Fig. 5**). The aim of this analysis was to compare the two methods (**Table 3**). In all cases, the HPTLC method presented in this work had a sensitivity comparable to that of the spectrophotometric analysis. However, our method allows for a broader range of lipid analysis than the sulfo-phospho vanillin assay, which is not lipid specific.

**Fig. 4:**
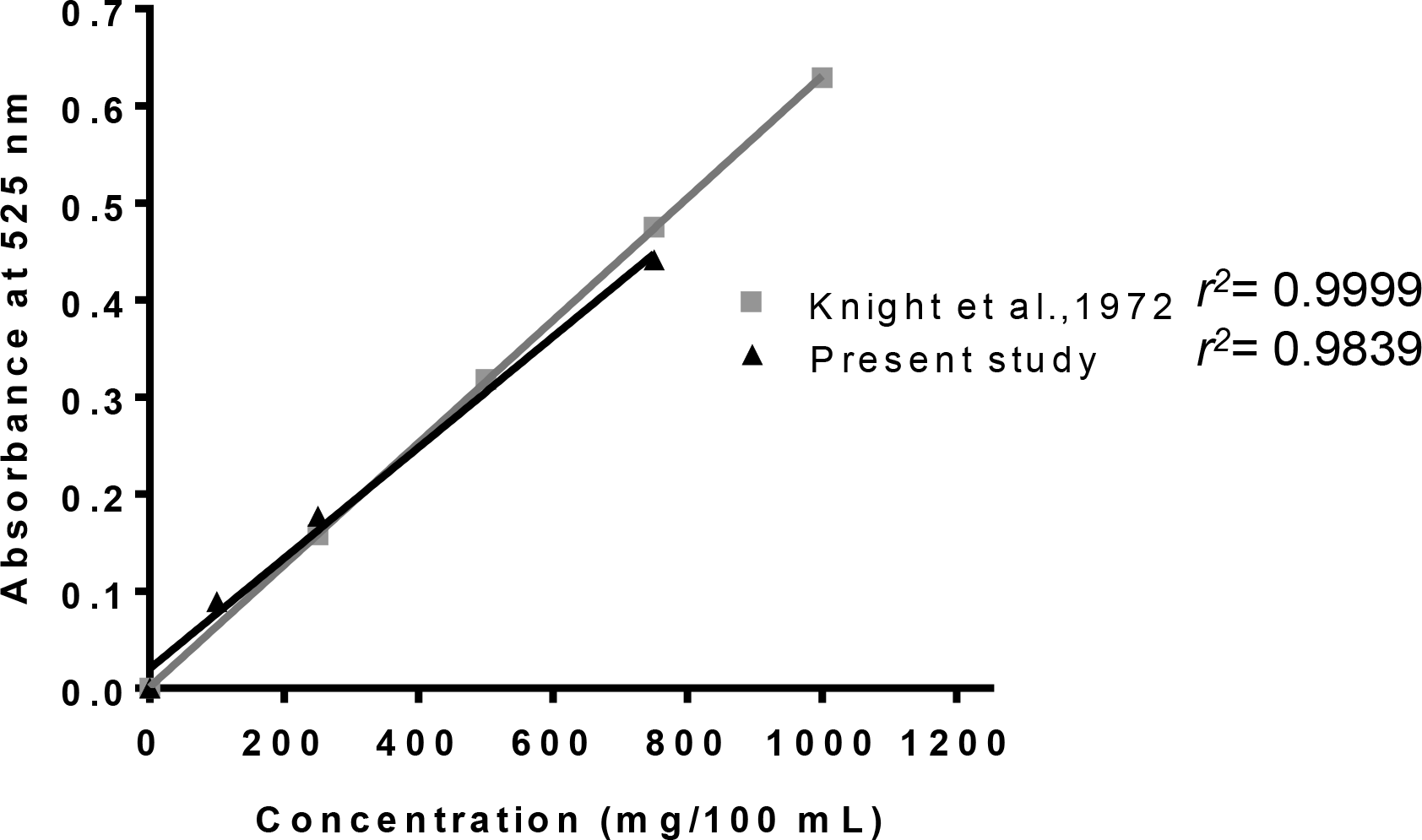
Comparison of the methods from Knight et al. [16] with the present one for the analysis of oleic acid. After spectrophotometric quantification of lipids using the phospho-vanillin method [16], standard curves obtained for the newly developed method and the phospho-vanillin method were compared for oleic acid.

**Fig. 5:**
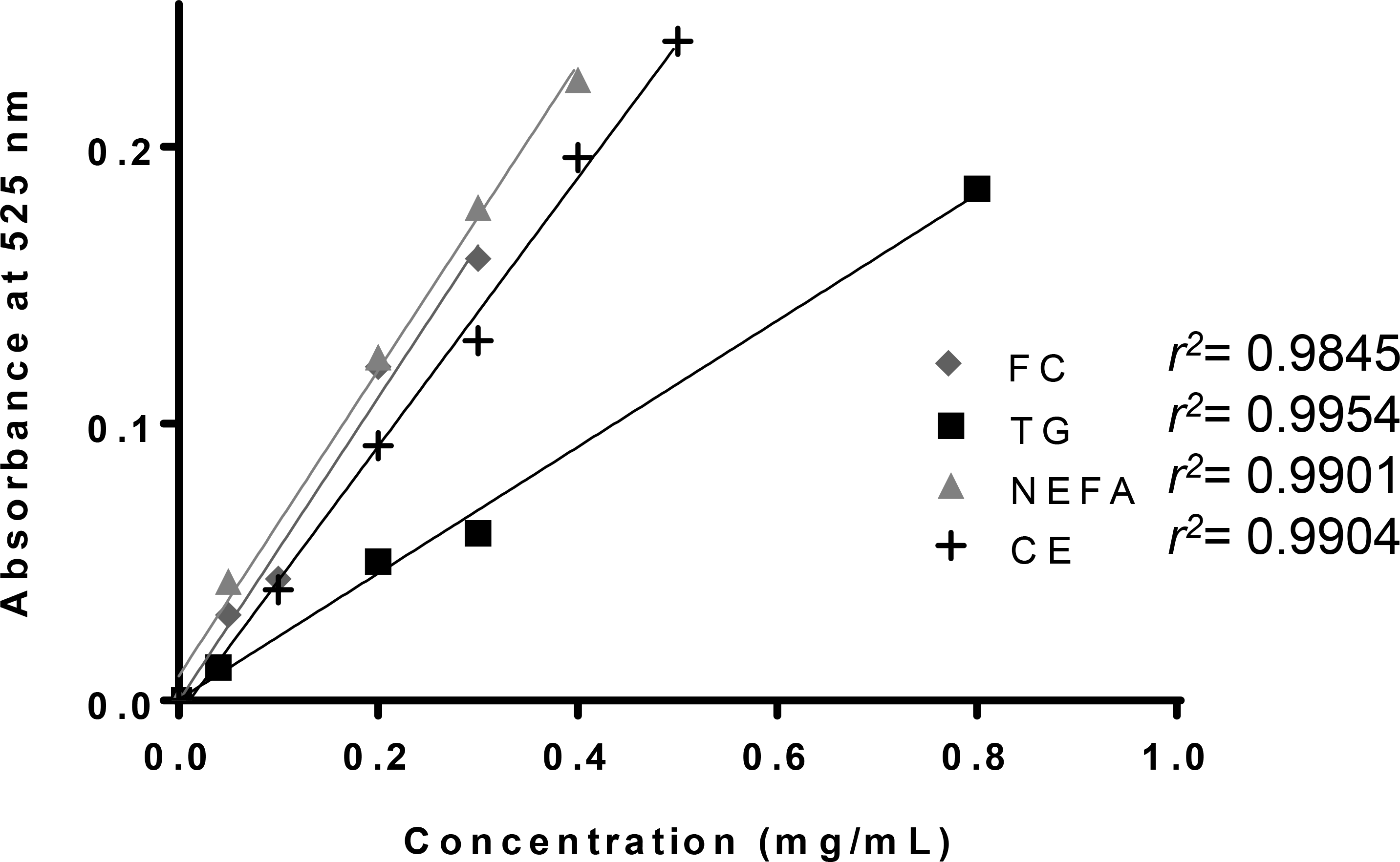
Quantification of the various lipids using the methods from Knight et al. After spectrophotometric quantification of lipids using the phospho-vanillin method [16], standard curves obtained with the phospho-vanillin method were plotted for free cholesterol, triglycerides, cholesteryl esters, and oleic acid. FC: free cholesterol, NEFA: non-esterified fatty acids, TG: triglycerides, CE: cholesteryl esters.

**Table 3:** Comparison of the lipid analyses performed by HPTLC and using the spectrophotometer method.

| HPTLC Method |  |  | Spectrophotometry method |  |  |
| --- | --- | --- | --- | --- | --- |
| Lipid | Concentration | Sensitivity | Lipid | Concentration | Sensitivity |
| TG | 0.04 mg/mL | 0.40 µg | TG | 0.04 mg/mL | 0.36 µg |
| NEFA | 0.05 mg/mL | 0,50 µg | NEFA | 0.05 mg/mL | 1.29 µg |
| FC | 0.05 mg/mL | 0,55 µg | FC | 0.05 mg/mL | 0.93 µg |
| CE | 0.04 mg/mL | 0,63 µg | CE | 0.04 mg/mL | 0,77 µg |

FC: free cholesterol, NEFA: non-esterified fatty acids, TG: triglycerides, CE: cholesteryl esters.

The accuracy of the method was also established using the recovery technique (i.e., external standard addition method). A known amount of standard was added at four different levels to a pre-analyzed sample. Each determination was performed three times. The results of the recovery study presented in **Table 4** were found to be within acceptable limits (value ±10%).

**Table 4:** Recovery studies of the different lipids after 50%, 100% and 150% addition of the standard lipid solution.

| Lipid | Amount at 100% (µg) | Levels | Expected amount to be assayed (µg) | Area observed (AU) | Amount calculated (µg) | Recovery (% mean ± SD) | n |
| --- | --- | --- | --- | --- | --- | --- | --- |
| FC | 4 | 50% | 2 | 65084 | 1.99 | 99.00 ± 1.02 | 3 |
| | | 100% | 4 | 130569 | 4.00 | 100.00 ± 0.59 | 3 |
| | | 150% | 6 | 193568 | 5.93 | 98.90 ± 0.72 | 3 |
| | | 200% | 8 | 253054 | 7.75 | 96.90 ± 0.58 | 3 |
| NEFA | 4 | 50% | 2 | 27574 | 1.98 | 99.00 ± 1.05 | 3 |
| | | 100% | 4 | 55833 | 4.00 | 100.00 ± 0.68 | 3 |
| | | 150% | 6 | 75504 | 5.41 | 90.00 ± 1.10 | 3 |
| | | 200% | 8 | 106589 | 7.63 | 95.00 ± 1.09 | 3 |
| TG | 4 | 50% | 2 | 25092 | 1.97 | 99.00 ± 0.57 | 3 |
| | | 100% | 4 | 50916 | 4.00 | 100.00 ± 0.88 | 3 |
| | | 150% | 6 | 68614 | 5.39 | 90.00 ± 0.96 | 3 |
| | | 200% | 8 | 99580 | 7.82 | 97.70 ± 0.90 | 3 |
| CE | 4 | 50% | 2 | 54984 | 1.82 | 91 ± 0.77 | 3 |
| | | 100% | 4 | 120823 | 4 | 100 ± 0.59 | 3 |
| | | 150% | 6 | 174019 | 5.76 | 96.00 ± 1.01 | 3 |
| | | 200% | 8 | 237512 | 7.86 | 98.28 ± 0.95 | 3 |

FC: free cholesterol, NEFA: non-esterified fatty acids, TG: triglycerides, CE: cholesteryl esters.

### Precision

Repeatability and intermediate precision were determined at different concentrations levels. The percent relative standard deviation (%RSD) values for repeatability and intermediate precision as shown in **Table 5** reveal that the proposed method is precise.

**Table 5:** Repeatability and intermediate precision data for the different lipids.

|  |  |  |  |  |
| --- | --- | --- | --- | --- |
| Quantity (µg) | **2** |  |  |  |
| Lipid | FC | NEFA | TG | CE |
| Repeatability (%) | 7.48 | 10.01 | 8.49 | 8.16 |
| Intermediate precision (%) | 7.49 | 13.60 | 11.60 | 10.97 |
| Quantity (µg) | **8** |  |  |  |
| Lipid | FC | NEFA | TG | CE |
| Repeatability (%) | 6.20 | 7.44 | 11.46 | 8.96 |
| Intermediate precision (%) | 6.80 | 7.99 | 13.59 | 9.88 |

FC: free cholesterol, NEFA: non-esterified fatty acids, TG: triglycerides, CE: cholesteryl esters. n = 10 for each concentration and each lipid.

### Limit of detection (LOD)and limit of quantification (LOQ)

Limit of detection (LOD) and limit of quantification (LOQ) values are presented in **Table 6**. This method is therefore sensitive for our analyses. Importantly, the LOQ values determined as described in the Material and Method section (**Table 6**) are comparable to those derived from the method using the Eurachem approach (**Fig. 6**) [15].

**Table 6:** Method validation parameters for the quantification of the different lipids.

| Lipid | FC | NEFA | TG | CE |
| --- | --- | --- | --- | --- |
| LOD (ng/spot) | 556 | 1746 | 1284 | 639 |
| LOQ (ng/spot) | 913 | 3092 | 2596 | 1593 |
| Regression coefficient | 0.9858 | 0.9991 | 0.9993 | 0.9987 |
| Cochran C ($C_{\text{calc}}/C_{\text{ref}}$, $p < 0.01$) | 0.4902/0.5702 | 0.4520/0.5702 | 0.3600/0.5702 | 0.3343/0.5702 |

FC: free cholesterol, NEFA: non-esterified fatty acids, TG: triglycerides, CE: cholesteryl esters.

**Fig. 6:**
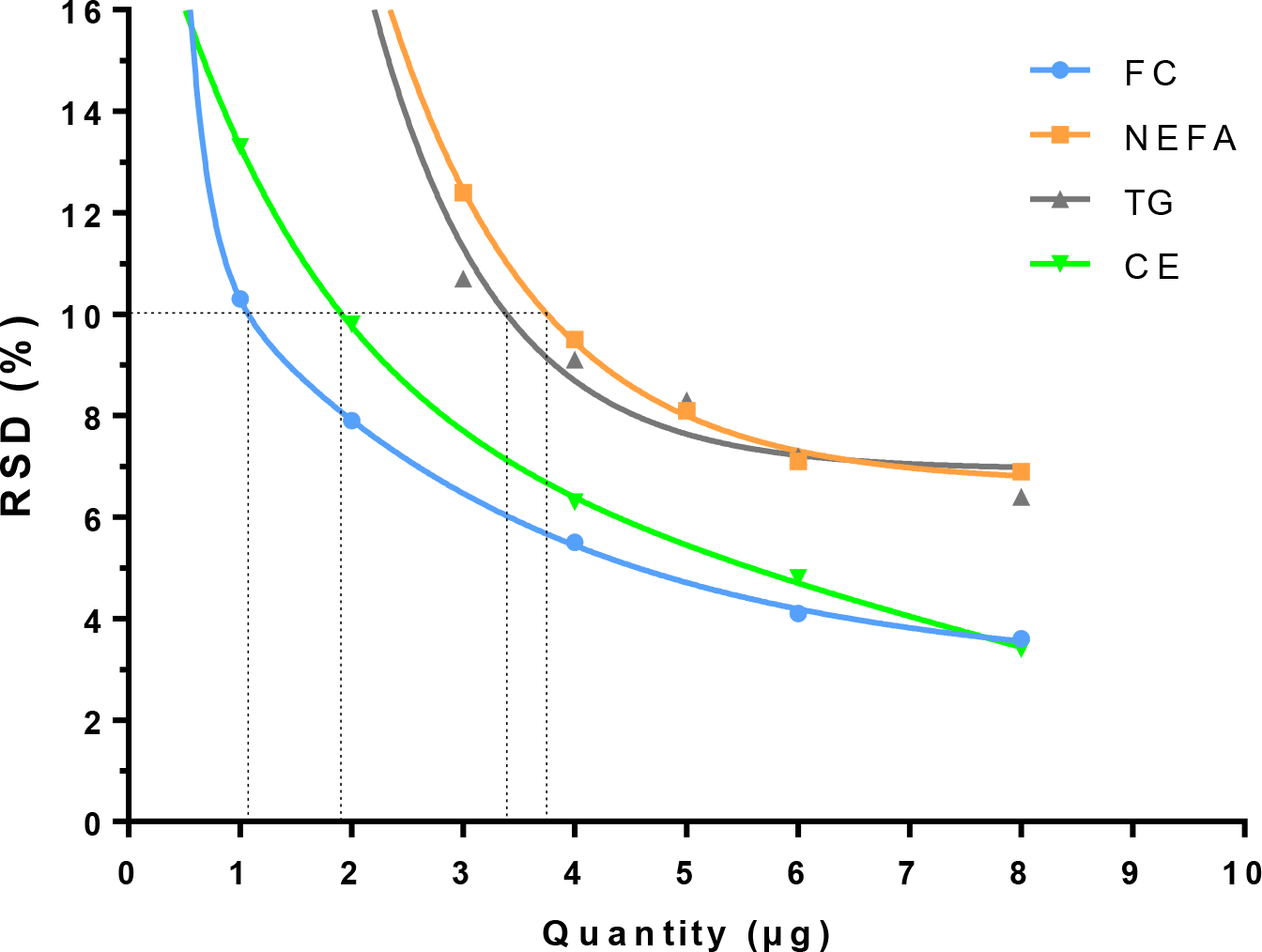
Precision profile obtained for the different lipids. RSD values were determined for analyses using 1-8μg of FC and CE or 2-8μg of NEFA and TG. LOQ values are indicated for an RSD of 10%.

### Specificity and Selectivity

Specificity of the method was assessed by analyzing authentic lipid standards. Identity of the lipids present in three different mammary epithelial cell lines (MCF-10A, MCF-7, and MDA-MB-231; after Bligh and Dyer extraction [13]) was determined by comparison of the lipids’ Rf with values obtained for authentic lipid standards (**Fig. 7**). All these cell lines contain triglycerides, non-esterified fatty acids, cholesteryl esters, and free cholesterol. This assay could allow us to determine the effect of a specific treatment on lipid metabolism (non-esterified fatty acids, free cholesterol, cholesteryl esters, triglycerides) using these cell lines. Interestingly, the MCF-7 cell line contains significantly less EC compared to the non cancerous MCF-10A cell line and the more aggressive MDA-MB-231 cell lines. FC, NEFA, and TG levels were not significantly different. The significance of the increased EC content observed in MCF-10A and MDA-MB231 cells remains to be determined. Importantly, these findings were confirmed by fluorescent microscopy using the neutral lipid stain, Nile Red (**Fig. 8**).

**Fig. 7:**
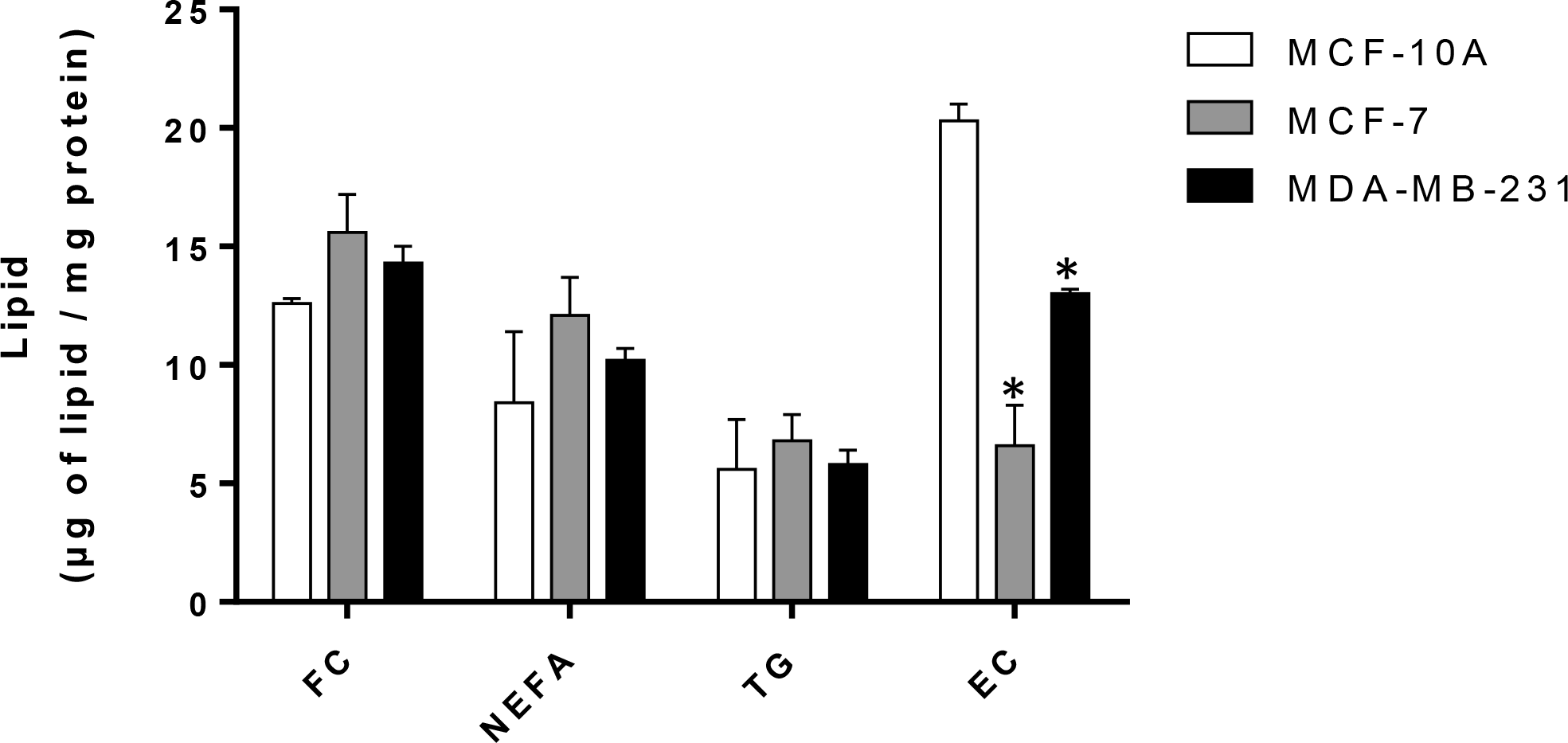
Analysis of a lipid extracts obtained from mammary cell lines. Cells were grown in DMEM 10% FBS. 10^6^ cells were collected and lipids were extracted by the method of Bligh and Dyer [13]. After extraction, lipids were separated by HPTLC as described in the Material and Methods section. After quantification, the composition of each cell line for the four different lipids was reported per mg of cell proteins.

**Fig. 8:**
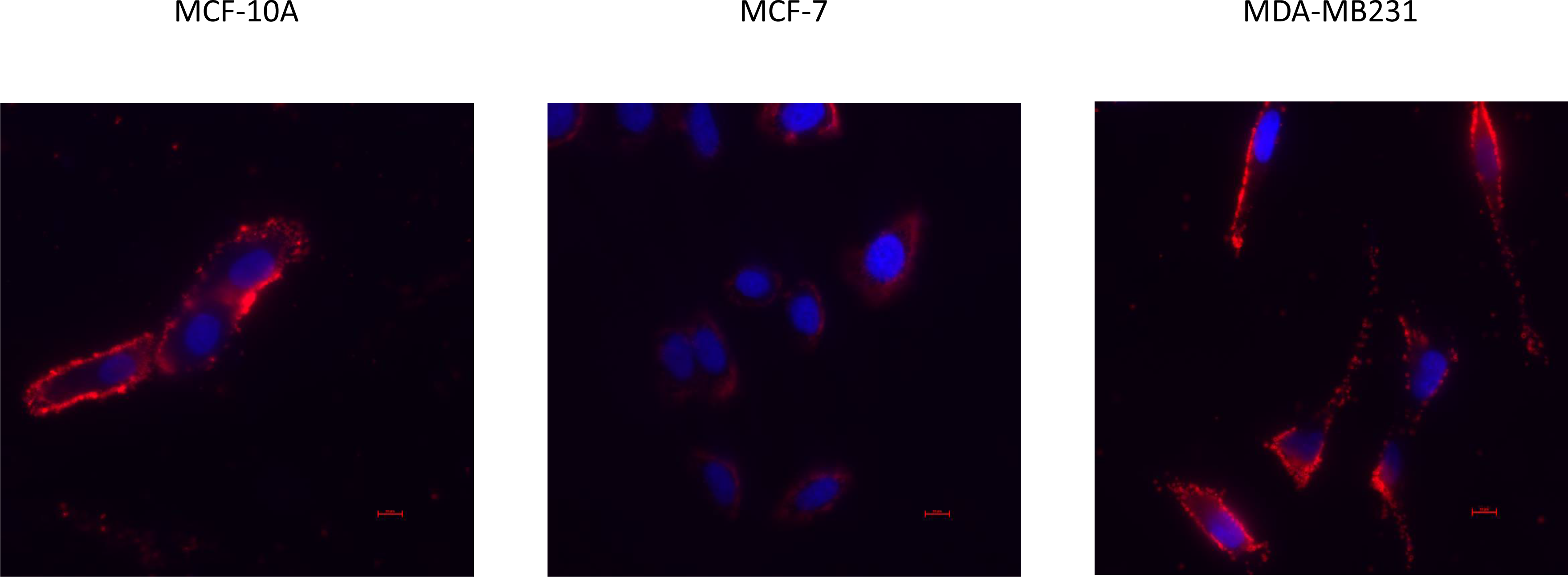
Analysis of a neutral lipid content in human mammary cell lines by fluorescence microscopy using Nile Red. Cells were seeded on coverslips and incubated in complete media. After 1 day in culture, cells were fixed in 2% paraformaldehyde, incubated with 1% BSA in PBS, and incubated with 300 nM Nile red (in PBS containing calcium and magnesium (CM)). After several wash in PBS-CM cells were mounted using SlowFade® diamond antifade reagent containing DAPI. Cells were observed on a Nikon TI-S microscope and analyzed using both NIS-BR software (Nikon, France). Red: Nile red staining (neutral lipids); blue: DAPI staining (nuclear stain).

Finally, other related lipid species did not have the same Rf as those studied in the present work. Diacylglycerol and cardiolipin had Rf of 0.28 and 0.01, respectively. Importantly, all types of phospholipid remain at the origin.

### Statistical analysis

For these lipid determinations, variance homogeneity of our data standard curve was verified using the Cochran’s C test for homoscedasticity (**Table 6**). Furthermore, to validate the linear regression model and test if the slope was different from 0, a Fischer’s test was applied. The value obtained for each lipid type was compared with the values from the F table. For this study, the test was accepted for all lipids at *p*<0.05. Our accuracy results show that the values obtained by HPTLC are significantly correlated with the amounts of lipids used in each case (**Table 6**). It therefore validates the use of our method for the quantification of these lipids in plasma, tissue, or cell extracts.

### Conclusions

The present study demonstrates that our HPTLC-densitometric analysis method is reproducible, repeatable, selective, and accurate for the analysis of the different lipids examined. Moreover, this technique is simple, fast, robust, and less expensive than HPLC. Multiple detection methods can be used in this context: UV (254 or 366 nm) or white light after derivatization using various reagents. However, we found that the system used in the present work gave the best results. Another important advantage is that several samples can be analyzed at once, thereby limiting the use of solvent and accelerating the analyses. Moreover, this method may allow us to quantify free cholesterol, cholesteryl esters, non-esterified fatty acids, and triglycerides present in plasma, adipose tissues, or other tissue/cell extracts containing these types of lipids. As an example, we show that this HPTLC-densitometry method can be routinely used for the quantification of these lipid classes in cancer cells.

## Acknowledgements

This work was supported by the Faculty of Medicine at the University of Tours. JLGP was supported by Le Studium (Région Centre-Val de Loire, France). PGF was supported by grants from INCa PLBio (2018-145), the Lipids ARD2020-Biodrug project (Région Centre-Val de Loire, France), La Ligue contre le Cancer (Indre et Loire, Loir et Cher, and Vienne), by an Academic Research Grant from the Région Centre-Val de Loire (France) and by the Canceropole Grand-Ouest (Mature project). We thank Dr. Jacques Potier, Violetta Guerin, and Simon Bergerard for technical support.

## Conflicts of Interest

The authors declare that they have no conflict of interest.

